# The Automated Optimisation of a Coarse-Grained Force Field Using Free Energy Data

**DOI:** 10.1101/2020.08.13.250233

**Authors:** Javier Caceres-Delpiano, Lee-Ping Wang, Jonathan W. Essex

## Abstract

Atomistic models provide a detailed representation of molecular systems, but are sometimes inadequate for simulations of large systems over long timescales. Coarse-grained models enable accelerated simulations by reducing the number of degrees of freedom, at the cost of reduced accuracy. New optimisation processes to parameterise these models could improve their quality and range of applicability. We present an automated approach for the optimisation of coarse-grained force fields, by reproducing free energy data derived from atomistic molecular simulations. To illustrate the approach, we implemented hydration free energy gradients as a new target for force field optimisation in ForceBalance and applied it successfully to optimise the un-charged side-chains and the protein backbone in the SIRAH protein coarse-grain force field. The optimised parameters closely reproduced hydration free energies of atomistic models and gave improved agreement with experiment.

## Introduction

Computational tools have become very important in revealing the driving forces in bio-molecular processes; in particular, molecular dynamics (MD) simulations provide a physically motivated picture based on Newton’s equations of motion coupled with empirical model potentials (force fields) ^1^. Classical atomistic (AT) models provide a detailed representation of the system with computational effort that scale as O(N log N) with the number of atoms, but are inadequate for simulations of very large systems over long timescales^2^. Coarse-grained (CG) models currently represent one of the most important approximations for the construction and simulation of larger systems^3,4^. By subsuming groups of atoms into single interaction sites, much faster calculations can be realised. However, a disadvantage of CG models is the loss of accuracy associated with reducing the number of interacting particles. Moreover, coarse-graining typically smooths the energy landscape compared to classical atomistic models, diminishing the energy barriers between different states and reducing trapping in energy minima^5^. This can greatly affect calculated thermodynamic properties such as equilibrium structures and dynamic properties such as the rates of conformational changes. Despite these drawbacks, CG models have become a widely used approximation, allowing us to extend spatial and temporal scales for the simulation of bigger and more complex systems. Given this, new approaches for the optimisation of CG models are highly desirable.

The accuracy of a force field depends in part on the empirical parameters in the model, which are usually determined by fitting simulation results to a training data set (i.e. the *targets*). For example, these targets can come from supermolecule calculations such as QM simulations or experimental information, but often such data are not available for the system of interest. All these difficulties, in conjunction with its iterative nature and complexity, mean that force field optimisation is something of a black art^6^. Different frameworks and approximations to optimise parameters have been proposed: 1) *ad-hoc* methods where parameters are iteratively adjusted until a specific property can be reproduced or stable simulations achieved ^7-9^, 2) machine learning methods that have been used in tandem with QM calculations^10,11^, and 3) force or energy matching to reproduce QM calculations or other simulation data^6,12,13^.

ForceBalance^14,15^ is an automated parameter optimisation method and software package that enables reproducible development of force field parameters. It has been used for the optimisation of different types of force fields, such as a series of water models (iAMOEBA^16^, AMOEBA14^17^, TIP3P-FB, TIP4P-FB^15^ and uAMOEBA^18^), a united-atom phospholipid bilayer model (gb-fb15)^19^, and an all-atom protein force field (AMBER-FB15)^20^. ForceBalance is able to incorporate multiple sources of experimental or simulated reference data. The objective function to be minimised in parameter space is a weighted sum of squared differences between the reference and calculated properties, with a regularisation term added that penalises large parameter deviations from their initial values to prevent overfitting. A harmonic penalty function, which corresponds to a Gaussian prior distribution, is usually used. ForceBalance uses a trust-radius Newton-Raphson optimiser that can efficiently optimise the objective function to within the statistical noise of the simulation after 5-10 iterations; other gradient-based and stochastic optimisation procedures may also be used in a modular fashion (e.g. L-BFGS, Simplex and Powell algorithms). The physical force field parameters are mapped to abstract optimisation variables of order one to improve the conditioning of the optimisation problem – this also enables one to adjust the regularisation strengths applied to different parameter types. The molecular mechanics property calculations are automated by interfaces to classical molecular dynamics software packages (*engines*) such as GROMACS^21^, TINKER^22^ and OpenMM^23^. Properties previously used in ForceBalance range from energies, atomistic forces, and vibrational modes from *ab initio* calculations^20^, *ab initio* gas phase properties such as cluster interaction energies, temperature and pressure dependent bulk phase properties of liquids such as density, enthalpy of vaporisation, dielectric constant, thermal expansion coefficient, isothermal compressibility and isobaric heat capacity^15,17^, and lipid membrane properties such as area per lipid and deuterium order parameters^19^.

Hydration free energies (HFEs) are an important property for aqueous systems such as proteins. They help us to understand biological processes such as ligand recognition, protein-protein interactions, folding and conformational changes. Moreover, hydration free energies have been used for the validation of molecular force fields, and they are an integral part of the calculation and estimation of solubilities, partition coefficients and solute-solvent interactions^24-27^. For these reasons, use of solvation free energies as a parameterisation target for coarse-grained models may improve their performance. Moreover, it has been recently stated that there is considerable interest in methods that can automatically generate a coarse-grained model and are representative in terms of local structure and free energy changes^28^.

Here we present a general approach to optimise coarse-grain force fields by reproducing free energy gradients derived from atomistic simulations. We exemplify the method by optimising the SIRAH CG protein force field using atomistic hydration free energy (HFE) data in the ForceBalance software. The gradient of the hydration free energy is optimised to match the result from an AT simulation, with the goal of improving the CG solvation free energies as a consequence. The approach of fitting atomistic HFE gradients has the advantage of reducing the computational cost of the parameter optimisation because it does not require full HFE calculations of the CG model at every optimisation step. The parameters of charged and uncharged amino acids were both optimized, but we rejected the charged amino acid parameters because they failed validation tests. A full HFE calculation is carried out after CG model optimisation to validate the approach by comparison to atomistic and experimental HFEs. The newly optimised SIRAH-OBAFE (**O**ptimised **B**ased in **A**tomistic **F**ree **E**nergies) force field, is briefly evaluated in terms of conventional MD simulations of proteins in solution. To facilitate the development of the new force field we have also optimised the WT4 water model in SIRAH using experimental properties such as density, enthalpy of vaporisation and dielectric constant.

## Methods

### Optimisation based on free energy gradients: overview

To calculate the free energy difference between two states, X and Y, it is useful to include a coupling parameter to connect both states^29-32^. This coupling parameter, α, changes from 0 to 1, and can be expressed as a linear function of the potential energy U(**r**^N^; α) by

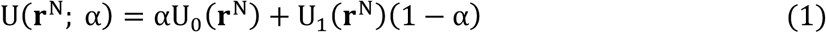

where **r**^N^ corresponds to the system coordinates of N particles, U_0_(**r**^N^) corresponds to the potential energy of a “reference system” and U_1_(**r**^N^) corresponds to the potential energy of a system of interest. α connects the two states through a physical or non-physical pathway. Based on thermodynamic integration theory^29^, one can express the difference in free energy between two states by:

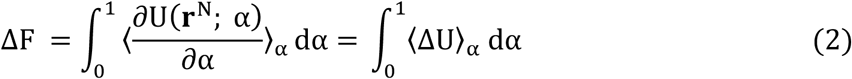

where the change in the free energy ΔF, between a reference state and a target state, can be computed from the integral between values of 0 (un-perturbed) and 1 (perturbed) of the ensemble average of the derivative of the potential energy with respect to the coupling parameter α. In the case of the linear coupling of U(**r**^N^; α), corresponding to equation (1), this is equivalent to the ensemble average of ΔU as a function of α, where ΔU is the internal energy change between the α = 0 and α = 1 states.

We have implemented a new mathematical expression for the optimisation of coarse-grained force field parameters based on free energy gradients from atomistic simulations. Starting with a set of simulations that evaluate <ΔU>_α_ for AT systems at selected values of α, we fit these values in our CG simulations by optimising the CG parameters, which indirectly improves the hydration free energies. The objective function that is minimized may be written as:

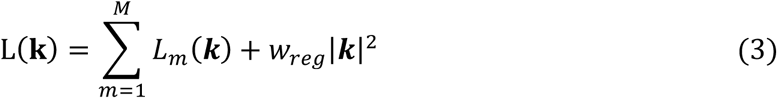

Here *L*_*m*_(***k***), called the target terms, are the contributions of each molecule to the objective function; in this work the parameters for each molecule are optimized separately, thus there is only one term in the sum. *L*_*m*_(***k***) is given by a weighted sum of squared differences between the AT and CG free energy gradients:

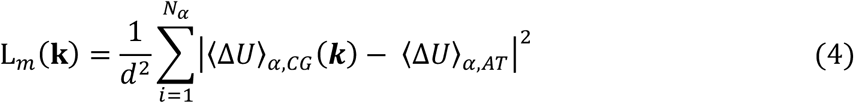

where ***k*** is the vector of dimensionless “mathematical” parameters being directly manipulated by the optimization algorithm, *L*(***k***) is the overall objective function, *L*_*m*_(***k***) is the contribution from molecule *m*, and *w*_*reg*_ is a regularization term, here set to 0.01 to ensure that large excursions in the parameters are properly penalized without being overly restrictive. The ***k***-vector is related to the physical force field parameters in the simulation ***K*** by a shifting and scaling as:

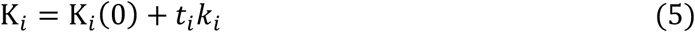

where *K*_*i*_ and *K*_*i*_(0) are the current and initial values of the force field parameter, and *t*_*i*_ is a scaling factor, also called the prior width, that carries the same dimension as *K*_*i*_ and represents the expected variation of the force field parameters over the course of the optimization.

In order to optimize the objective function efficiently, the first derivatives of the simulated quantities with respect to force field parameters are needed. The analytical derivative of <ΔU>_α_ with respect to the force field parameters can be obtained as:

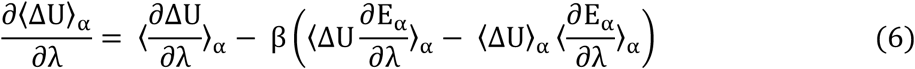

where λ corresponds to the force field parameter, <ΔU>_α_ is the ensemble average of the energy difference between α = 0.0 and α = 1.0, simulated at a defined α value, ΔU corresponds to the instantaneous energy difference for each snapshot between α = 0.0 and α = 1.0, and *E* is the potential energy of the system at α. Rather than optimising the free energies directly, we optimise against the ensemble average of the free energy gradients at specific α values, <ΔU>_α_. The derivative of the free energy gradients, <ΔU>_α_, with respect to the force field parameters λ is composed of ensemble averages of instantaneous ΔU values, and derivatives of ΔU and the potential energy with respect to the FF parameters, at each α point used, where both derivatives are obtained numerically by finite difference using snapshots from the corresponding trajectories.

### Optimisation of a CG protein force field: uncharged side-chains and backbone

A workflow showing the steps followed in this work, and separated into four main stages, is presented in figure 1. Briefly, hydration free energies for atomistic systems are calculated by decoupling both van der Waals and charge parameters. Then, atomistic free energy gradients are collected as an average of ΔU values, at simulations with different α values, <ΔU>_α_. These data are used to optimise each specific CG side-chain (or the backbone) with its corresponding <ΔU>_α_ value. Then, parameters corresponding to the smallest objective function are collected. These parameters are then used to re-calculate new hydration free energies of the CG side-chains. See supporting information for more details of the stages shown in figure 1.

**Figure 1.**
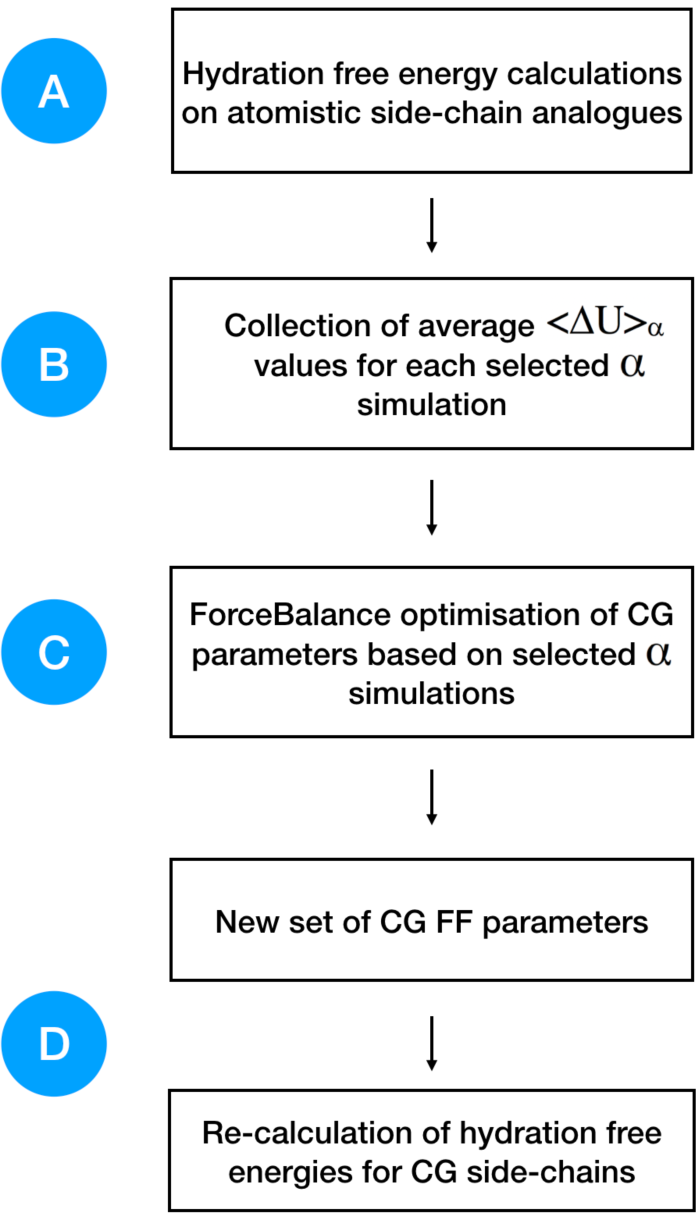
General workflow for the CG force field optimisation. Free energy gradients are collected from atomistic simulations and used as optimisation targets in ForceBalance. New parameters are obtained and later used in the re-calculation of hydration free energies for CG beads (side-chains and backbone). Letters from A to D correspond to each of the main stages in the optimisation and validation process (see SI).

### Hydration free energies of charged side-chains

The calculation of hydration free energies for charged systems is a more complex process compared to the classical use for uncharged systems. The standard raw hydration free energy 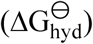 for an ion is calculated as the sum of three processes: charging (ΔG_chg_), cavitation (ΔG_cav_) and a standard convention term (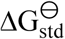, which is equal to 7.95 kJ·mol^-1^, considering a water density of 997 kg·m^-3^ at a pressure of 1 atm^33^), as:

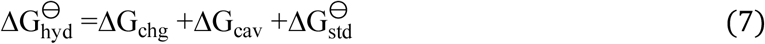

The cavitation term corresponds to the creation of a molecule in solution through the scaling of intermolecular Lennard-Jones interactions, coupled to a parameter α.

The calculation of raw charging free energies 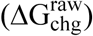 is especially sensitive to the chosen simulation methodology^33-35^ where different corrections have been introduced to alleviate these effects (see SI). Following these corrections^33-35^, raw hydration free energies and these corrections (ΔG_cor_, see SI) were used to calculate the methodology-independent free energy values for the charged side-chains, as:

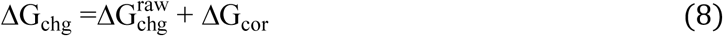

These corrections, and their application, have been demonstrated before for monoatomic^34,35^ and polyatomic ions^33^. They are usually named as type A, B, C and D corrections, which are related to approximations in the electrostatic interactions (A), approximations of the system size (finite) (B), deviations of the solvent generated electrostatic potential given the choice of an inappropriate summation scheme (C), and a wrong estimation of the dielectric constant for solvent model used (D), respectively. In the case of polyatomic ions (such as the charged side-chains used in this work), numerical solutions of the Poisson equation are needed to obtain an estimation of the charging free energy in an idealised system that obeys a macroscopic regime (non-periodic with Coulombic electrostatic interactions) and based on the experimental solvent permittivity 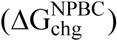. Simulations of a periodic systems with a specific electrostatic scheme and based on the model solvent permittivity are also needed (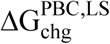 for a periodic boundary condition system using a **L**attice-**S**ummation scheme). These two terms can be used for the calculation of A+B+D corrections from continuum electrostatic calculations. See supporting information for more details.

A type C_1_ correction is required for lattice-summation (LS) and Barker-Watts reaction field (BM) schemes, and corrects the P-summation (atom-based cut-off) implied by these schemes to a proper M-summation (molecule-based cut-off). This correction is calculated analytically. See supporting information for more details.

Finally, and summarising all the necessary methodology-dependent corrections, standard hydration free energies were calculated as:

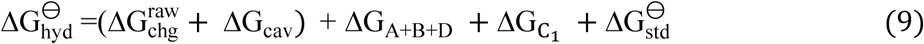

### Optimisation of charged side-chains

Optimisations for the charged side-chains were performed in a similar fashion as the case for uncharged side-chain and the protein backbone, where free energy gradients from atomistic simulation were used as optimisation target for the CG parameters (see equation 3). As this type of free energy calculation is methodology dependent, the inclusion of corrections is required. Assuming that the final free energies between the atomistic and coarse-grained models must be equal, the sum of their free energy gradients and the necessary correction gradients must be equal as well. Since the corrections are added *ex post*, the fitting data used is given as,

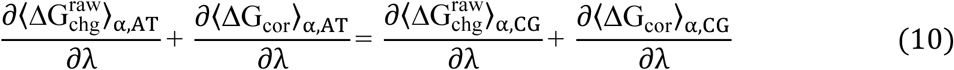

and moving the property that we want to optimise to one side,

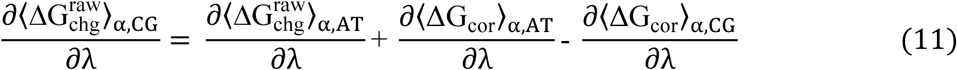

where 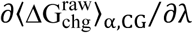 and 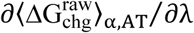 correspond to the derivative of the raw charging hydration free energy gradients with respect to the force field parameters (at a specific α value), for a coarse-grained and atomistic system, respectively. ∂⟨ΔG_cor_⟩_α,CG_/∂λ and ∂⟨ΔG_cor_⟩_α,AT_/∂λ are the derivatives of the free energy corrections with respect to the force field parameters (at a specific α value), for a coarse-grained and atomistic system, respectively. The derivatives of the corrections were calculated using finite differences, based on a set of α values between 0.4 and 1.0, where the parameters were scaled accordingly (i.e. for α=0.9, parameters were scaled to a 90% of their original value). See SI for more details on the calculation of these corrections and the protocol used in optimisation runs.

### The SIRAH model

Our new parameterisation approach has been applied to the optimisation of the SIRAH force field^36^, a CG force field and a promising alternative to conventional atomistic protein force fields. Unlike MARTINI^37^, SIRAH does not use elastic networks to overcome the problem of secondary structure stability. The use of a higher resolution backbone representation produces hydrogen bond-like interactions, which stabilise the secondary structure. Moreover, SIRAH models long-range electrostatic interactions using the particle mesh Ewald method (PME) and a dielectric constant of unity. At the moment, the SIRAH force field contains parameters for DNA^38^, water^39^, proteins^36^ and DMPC lipid^40^, and it has been used in the simulation of protein-DNA interactions^41^, hybrid AT/CG simulations^42^ and in the implementation of a supra-CG water model for the simulation of virus-like particles^43^.

The SIRAH CG protein model uses a higher resolution backbone compared to previous CG models, where positions for nitrogen, α-carbon and oxygen are maintained. Each bead possesses its own partial charge, which helps to stabilise secondary structures through the formation of hydrogen bond-like interactions. Dihedral angles define the secondary structure for the system, forcing the existence of the two main conformations, α-helices and β-strands. Side-chains are modelled using one to five pseudo-atoms and partial charges are placed based on the number of hydrogen-bond acceptors and/or donors. Van der Waals parameters were set using an ad-hoc procedure, and van der Waals interactions are calculated based on the Lorentz-Berthelot combining rules, with the addition of some corrections^36^. The SIRAH water model (WT4) is represented by four linked beads in a tetrahedral geometry, each with a specific partial charge. Each CG water molecule represents approximately 11 atomistic water molecules based on the mass of CG beads (50 au)^39^. A new, updated version of the SIRAH protein model was recently released, named as SIRAH 2.0, where corrections were made to bonded and non-bonded interactions of amino-acids, showing decreased RMSD values up-to 0.1 nm, for different protein systems, compared to the previous SIRAH 1.0 version^44^.

### Optimisation of the WT4 model

For the WT4 model optimisation, three condensed-phase properties were optimised: density, enthalpy of vaporisation and dielectric constant. Experimental values (taken from ref. ^15^) for these properties were used as targets, at 298.15 K and 1 atm. The trust-radius Newton-Raphson algorithm was used to minimise the objective function (see SI for more details). For this work, the optimisation was regularised using a Gaussian prior that is centred on the original SIRAH parameter. This is done to prevent the optimisation from changing the parameters too much and to avoid over-fitting, adding a penalty that is applied to the objective function. Conceptually speaking, addition of a penalty function is equivalent to imposing a prior probability distribution on the parameters. Only non-bonded parameters were optimised, including van der Waals sigma (σ) and epsilon (ε) values, and partial charges.

100 optimisation cycles were run, with the following simulation protocol: the system was minimised for 5000 steps using a steepest descent algorithm followed by an NPT equilibration time of 5 ns. Production runs were performed for 15 ns. A leap-frog algorithm was used to integrate Newton’s equations of motion with a time-step of 20 fs. Electrostatic interactions are calculated using the Particle mesh Ewald method^45^ with a direct cut-off of 1.2 nm and a grid spacing of 0.2 nm. A 1.2 nm cut-off was used for van der Waals interactions. The V-rescale thermostat^46^ and the Parrinello-Rahman barostat^47^ were used to maintain the temperature at 298.15 K and the pressure at 1 atm, respectively. The simulation protocol was based on the original publication of the SIRAH 1.0 protein force field^36^. All simulations were run with GROMACS v. 2018.2 ^48^. Statistical fluctuations in the thermodynamic properties dominated the objective function after 30 iterations, and the set of parameters with the lowest objective function was chosen as the best solution. Single point calculations were run three times, with the best parameter set, in order to estimate standard errors.

### Protein simulations

To briefly evaluate the optimised force field, a series of proteins with sizes ranging from 585 to 69 residues were simulated (most were proteins tested in the original SIRAH 1.0 publication^36^). Coarse-grained molecular dynamics simulations were performed using the SIRAH 1.0/WT4, SIRAH 2.0/WT4 and the ForceBalance reparameterised SIRAH-OBAFE/WT4-FB force fields, for all the previously mentioned protein systems. Energy minimisation was carried out for 10000 iterations of the steepest descent algorithm. This was followed by an NPT equilibration dynamics procedure of 20 ns with positional restraints of 1000 kJ·mol^-1^·nm^-2^ applied to all the protein beads. Production runs were performed for 3 μs for each system with an integration time-step of 20 fs. Electrostatic interactions were calculated using the Particle Mesh Ewald procedure^45^ with a direct cut-off of 1.2 nm and a grid spacing of 0.2 nm. Non-bonded interactions were modelled using the Lennard-Jones potential with a cut-off of 1.2 nm. All simulations were run at 1 bar with the Parrinello-Rahman barostat^47^ and at 298.15 K with the v-rescale thermostat^46^. Systems were neutralised by adding Na^+^ and Cl^-^ ions up to a concentration of 150 mM. Root mean square fluctuations (RMSF) and root mean square deviations (RMSD) time series were calculated with GROMACS v.2018.2^48^.

## Results and Discussion

One of the main points that encouraged the development and improvement of these CG models, and also an important limitation of the SIRAH force field, is the inaccuracy of the hydration free energies of amino acid side-chains, which could limit its predictive power in protein simulations. Calculations of SIRAH 1.0 decoupling hydration free energies yield completely different results compared to all-atom OPLS-AA results, with mean unsigned errors against experiment (MUE) of 5.03 kcal·mol^-1^ vs. 1.04 kcal·mol^-1^, for SIRAH 1.0 and all-atom systems, respectively, calculated against experimental values (see below).

### Optimisation of the WT4 water model

We start our ForceBalance calculation with the optimisation of the WT4 water model, where only non-bonded parameters were optimised (charges, sigma and epsilon values). Three condensed-phase properties for liquid water were used as reference data: density, enthalpy of vaporisation and dielectric constant at 298 K and 1 atm. The original WT4 model is able to reproduce experimental thermodynamic properties such as the water density at 298 K, but it is less satisfactory in the prediction of other properties (i.e. dielectric constant, expansion coefficient, surface tension, etc.)^39^. In contrast, the new WT4 model (now called WT4-FB) overcomes the previous issue with the dielectric constant in the original model by accurately reproducing experimental values for the three properties together (table 1). Calculations of the thermal expansion coefficient yield similar results to those of the original model (11.8×10^−4^ K^-1^ vs. 11.6×10^−4^ K^-1^)^39^. Thus, optimising WT4 with ForceBalance does not necessarily improve all properties; the level of accuracy obtainable is dependent on the granularity of the CG representation and the choice of force field functional form.

**Table 1.** Comparison of WT4 and WT4-FB models against experimental water properties at 298 K and 1 atm^a^

| Property<sup>a</sup> | Expt. | WT4 | WT4-FB (this work)<sup>b</sup> |
| --- | --- | --- | --- |
| $\rho$ (kg·m<sup>-3</sup>) | 997.045 | 996.6 ± 0.3 | 995.4 ± 1.5 |
| $\Delta H_{\text{vap}}$ (kJ·mol<sup>-1</sup>) | 43.989 | 39.8 ± 0.2 | 43.7 ± 0.2 |
| $\epsilon_r$ | 78.409 | 123.7 ± 14.2 | 74.2 ± 12.3 |
| $\alpha$ (10<sup>-4</sup> K<sup>-1</sup>) | 2.572 | 11.6 ± 2.4 | 11.8 ± 2.7 |

<sup>a</sup> The calculated properties correspond to density ( $\rho$ ), enthalpy of vaporisation ( $\Delta H_{\text{vap}}$ ), dielectric constant ( $\epsilon_r$ ) and expansion coefficient ( $\alpha$ ), and the experimental data were obtained from reference 14.
<sup>b</sup> Full set of parameters for the WT4-FB model provided in table S2. Error are reported as standard errors based on 3 simulations.

### Optimisation of the SIRAH protein force field: uncharged side-chains and backbone

Our new approach for CG FF optimisation is based on using derivatives of the free energy gradients i.e. <ΔU>_α,_ at different values of the coupling parameter α, with respect to the force field parameters. We choose to work with free energy gradients due to their linear relationship with the easily computed “vertical energy gap”, <ΔU>_α_. In practice, the thermally averaged CG <ΔU>_α_, is fitted to atomistic <ΔU>_α_, where one or more selected values of the coupling parameter α are used to carry out the simulations.

HFEs were computed separately from the optimisation process. 10 sets of CG parameters were optimised representing 13 uncharged side-chains because of the shared mapping scheme and bead types for some groups of side chains; e.g. Asn/Gln share the same mapping, as do Ser/Thr and Val/Leu/Ile, and the backbone. Figure 2 and table S3 summarises the performance of our new set of parameters for uncharged side-chains and the backbone, now called SIRAH-OBAFE, together with the new WT4-FB force field, against HFEs from atomistic force fields (OPLS-AA^49^ for side-chains and AMBER-14SB^50^ for the backbone), the original SIRAH 1.0 force field^36^, the updated SIRAH 2.0 force field^44^, and experimental data^15^. Those atomistic force fields with optimum published reference data were chosen to be part of the parametrisation process; the CG force field should be agnostic to the all atom data from which it is parameterised.

**Figure 2.**
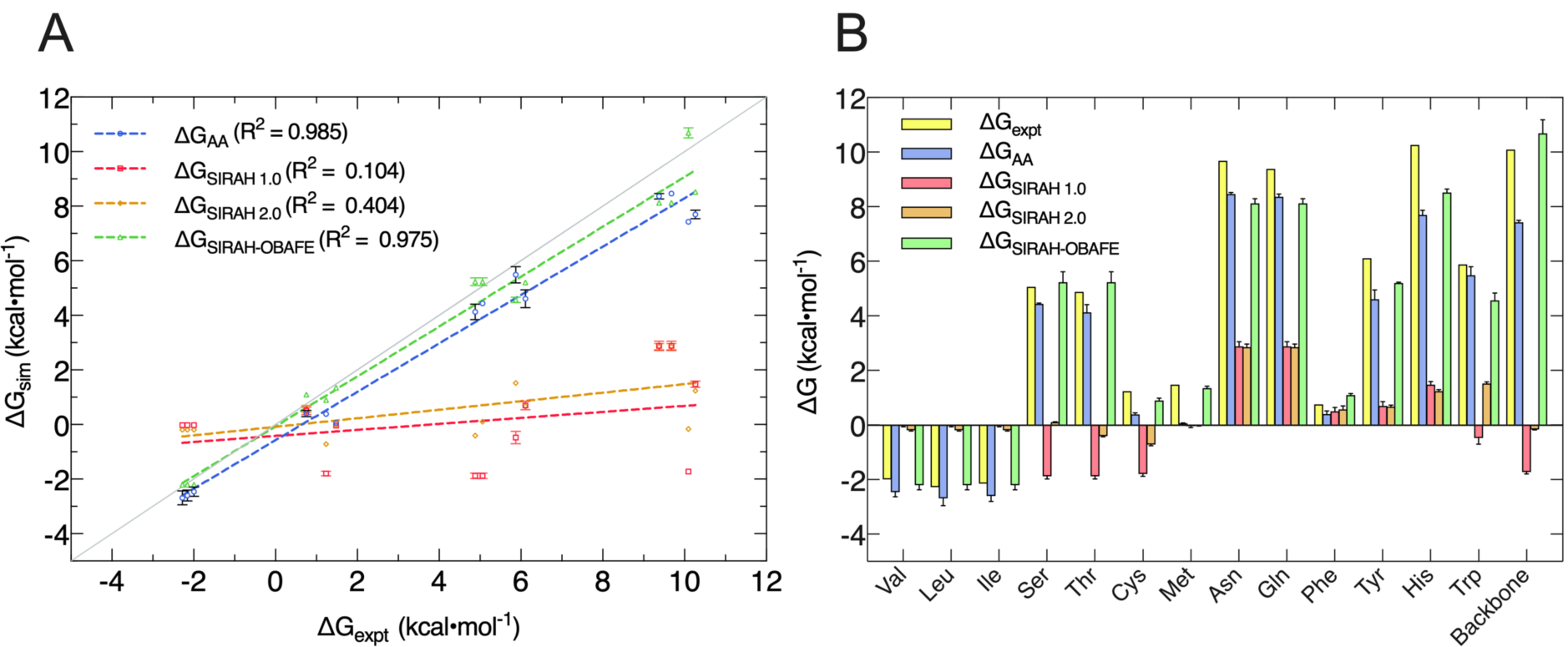
Comparison of decoupling HFEs from the new set of optimised parameters (SIRAH-OBAFE) against atomistic simulations (AA), the original SIRAH 1.0 force field, the latest version SIRAH 2.0 and experimental data. (A) Linear regression of predicted ΔG values for AA (blue), SIRAH 1.0 (red), SIRAH 2.0 (orange) and SIRAH-OBAFE (green) force fields, against experimental data. Each point represents a specific side-chain. The grey line represents a perfect fit (y=x), and R^2^ values are given in the inset legends. (B) Bar plot comparison of predicted ΔG values for AA (OPLS and AMBER-14SB) (blue), SIRAH 1.0 (red), SIRAH 2.0 (orange) and SIRAH-OBAFE (green) against experimental data (yellow; y axis) for all the neutral side-chains. Error estimates were calculated as standard errors based on three repeat simulations. For some cases, red bars appear to be missing as they are too small to be seen on the scale of the plot.

As can be seen, the original set of parameters in SIRAH 1.0 do not perform well for the prediction of decoupling HFEs, with an R^2^ of 0.104 against experimental values (Fig. 2A). A similar case is observed for the latest version SIRAH 2.0, with an R^2^ of 0.404 (Fig. 2A). SIRAH-OBAFE is able to greatly improve the agreement with experimental HFEs to be as good as atomistic force fields, with an R^2^ of 0.982 and 0.975, respectively. SIRAH-OBAFE reproduces the correct sign of several neutral side-chains where the previous SIRAH 1.0 model predicted the wrong sign, such as Ser, Thr, Cys and Trp (Fig. 2B). Significant improvements have been made to the HFEs of hydrophobic residues such as Val, Leu, and Ile; these share the same representation in SIRAH, using just one bead. The original SIRAH 1.0 and the updated SIRAH 2.0 models predict -0.02 ± 0.01 and -0.18 ± 0.01 kcal·mol^-1^ for the HFE, respectively (table S3), whereas SIRAH-OBAFE achieves a value of -2.26 ± 0.03 kcal·mol^-1^ (table S3); the latter value is much closer to OPLS-AA simulations and experiment which provide HFEs of (−2.45, -2.69, -2.59) and (−1.99, -2.28, -2.15) kcal·mol^-1^, for Val, Leu, and Ile, respectively (table S3). In the case of methionine, SIRAH-OBAFE produced even more accurate HFE values than the OPLS-AA model that provided the HFE gradients to which the CG model was fitted; we think this result is fortuitous and the differences are within the residual errors of the CG model vs. the AT reference (see SI related to the methionine case and figures S2 and S3).

### Optimisation of the SIRAH protein force field: charged side-chains

A different approach, compared to the optimisation of uncharged side-chains and backbone, was followed for the charged side-chains. We started with ForceBalance optimisation procedures, where the gradients of the raw charging free energies plus the gradient of the methodology-dependent corrections were used (see methods). Most of the ForceBalance optimisation results yield good agreement with experimental and AT hydration free energies (table 2, denoted as HFE-fitted), but the parameters were driven to unphysical values (see Fig. S4 and S5, for charge and Lennard Jones parameters, respectively). Given this, we conclude that our optimisation procedure works, but given the existence of few parameters to represent charged side-chains in SIRAH, and despite the use of regularization, over-fitting might be an unavoidable consequence in this case. Moreover, coarse-graining is an important simplification of the physics, where the option to fully reproduce complex properties, such as the free energy of charged entities, might not be possible. We have therefore decided to use the original SIRAH 1.0 parameters for charged side-chains, in combination with the hydration free-energy optimised parameters for backbone and uncharged side-chains, for the test of the SIRAH-OBAFE force field on protein systems.

**Table 2.**
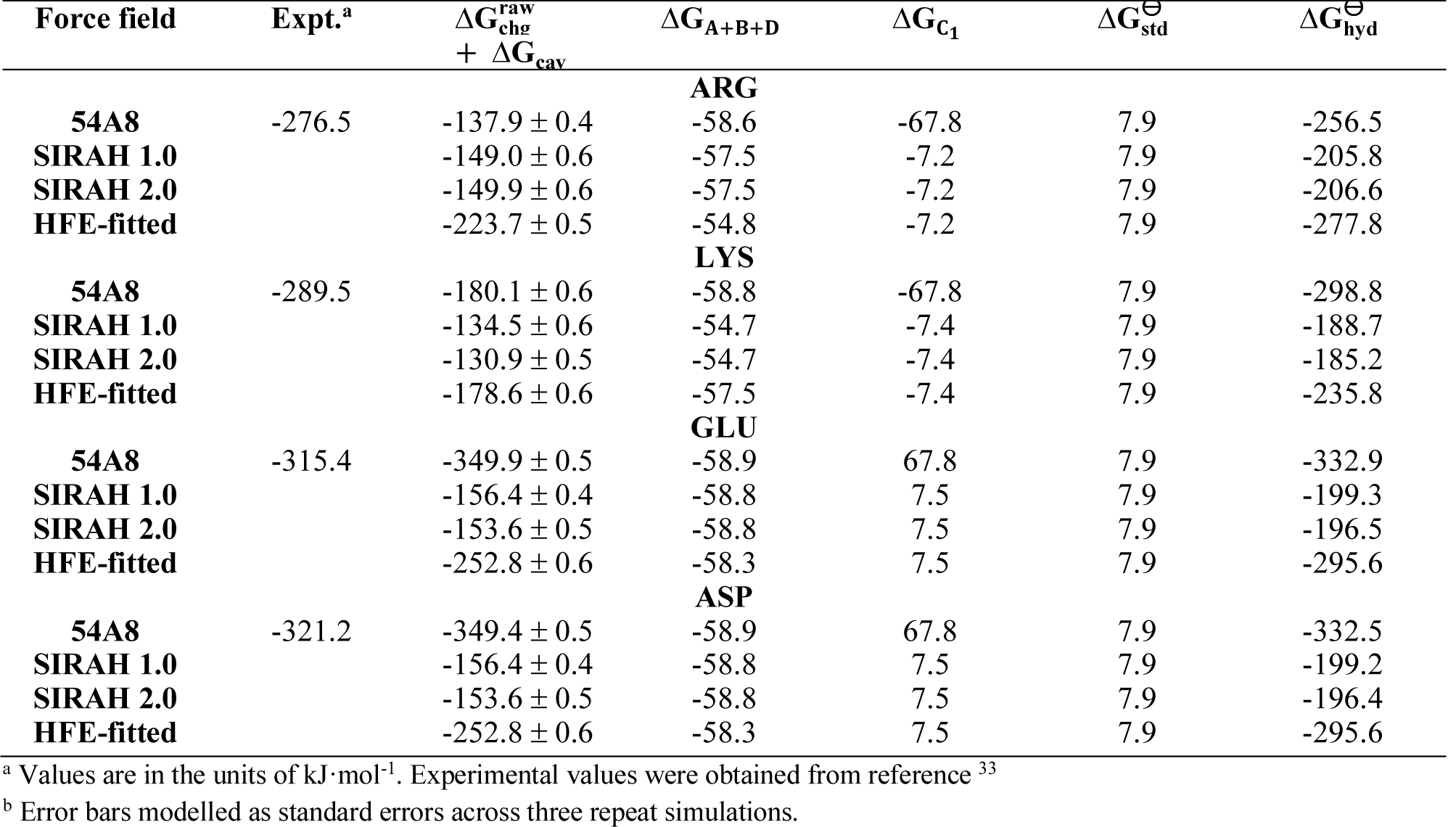
Hydration free energies of charged side-chains using the GROMOS 54A8, SIRAH 1.0, and SIRAH 2.0 force fields. HFE-fitted values are also included with the sole intention of comparison and discussion.

### Protein simulations

To test the performance of SIRAH-OBAFE in protein simulations, Cα RMSD analyses (with respect to the crystal structure) were performed on 6 protein system of different sizes. Simulations using the optimised SIRAH-OBAFE with the optimised WT4-FB were run for 3 μs. While the computed RMSDs are generally larger compared to atomistic simulations, all the simulations that used the optimised SIRAH-OBAFE model show improvements in protein stability with lower RMSD values throughout the whole trajectory with respect to the original SIRAH 1.0 and the updated SIRAH 2.0 force fields (Fig. 3). Even though the overall behaviour of the optimised SIRAH-OBAFE FF does not yield identical results compared to atomistic RMSDs, it shows an important improvement compared to the original SIRAH FF. As a simple comparison, atomistic simulation of systems with PDB codes 1QYO, 1RA4 and 1R69 (chosen from the original SIRAH 1.0 publication^36^) were run for 200 ns, showing average RMSD values of 0.270 nm, 0.06 nm and 0.148 nm, respectively. The original SIRAH 1.0 force field shows averaged RMSD values of 0.723 nm, 0.755 nm and 0.804 nm, while our optimised SIRAH-OBAFE force field shows averaged values (over the entire simulation) of 0.453 nm, 0.491 nm and 0.635 nm, for the same three cases, 1QYO, 1RA4 and 1R69, respectively. In the case of the updated SIRAH 2.0 force field, the overall behaviour of the RMSD timeseries is similar to the optimised SIRAH-OBAFE (Fig. 3), except for two larger systems, with average values (over the entire simulation) of 0.543 nm vs. 0.429 nm for 1E7I, and 0.601 nm vs. 0.453 nm for 1QYO (Fig. 3A and 3B). Even though the new RMSD values are not close to the atomistic RMSD and we would not necessarily expect them to be, there is as an improvement in the stability of protein systems based on our new optimisation approach. In more detail, calculating RMSD values against the last frame of the trajectories can give an insight whether the large RMSDs are due to large fluctuations or a change in conformation to a rigid conformer. This was performed using the updated SIRAH 2.0 and the optimised SIRAH-OBAFE force fields. Figure S6 summarise the results. As can be seen in the case of the 1E7I system, a big change of conformation is seen at around 1µs, which stabilise afterwards. In the other systems, it seems that the higher RMSD values observed in figure 4 are due to conformational drift across the simulation, with similar behaviours for both the updated SIRAH 2.0 and the optimised SIRAH-OBAFE force fields. RMSD fluctuations within a particular protein conformation are of the order of 0.2 nm.

**Figure 3.**
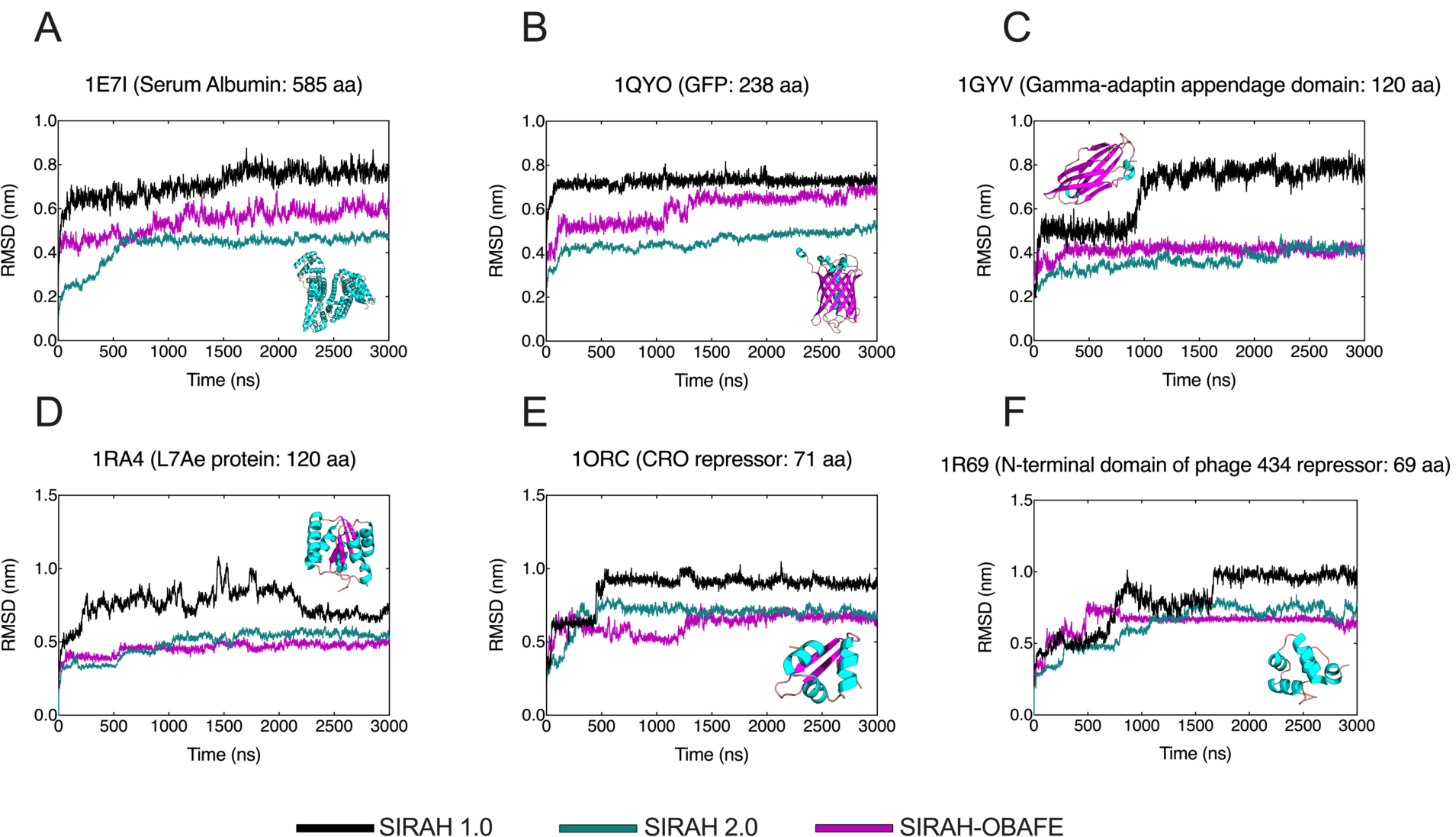
RMSD time series comparison between the original SIRAH 1.0 FF (purple), the updated SIRAH 2.0 FF (black) and the optimised SIRAH-OBAFE FF (green). RMSD trajectory analysis is shown as a time series comparison with respect to the Cα carbons of the CG representation to the crystal structure for (A) Serum albumin, (B) GFP protein, (C) Gamma-adaptin domain, (D) L7Ae protein, (E) CRO repressor and (F) the N-terminal domain of phage 434 repressor. PDB codes are shown in the figure titles and legend colours are shown at the bottom of the figure. Protein structures, corresponding to each of the simulated cases, are shown inside each plot. All simulations and analysis were run in GROMACS v.2018.2.

## Conclusions

In this work we propose a new and promising approach for parametrising coarse-grain force fields by optimising the CG force field parameters against free energy gradients derived from atomistic simulations. Our implementation of this method into ForceBalance enables full automation of the complex optimisation procedure and the incorporation of flexible choices of target data. It has been stated that there is considerable interest in methods that can automatically generate a coarse-grained model, and that are representative in terms of local structure and free energy changes. Our method paves the way to new optimisation procedures that rely on the use of free energy data as a target.

Non-bonded interaction parameters of un-charged side-chains and the backbone of the SIRAH coarse grain protein force field were optimised against hydration free energy gradients of atomistic simulation models, and compared against experimental hydration free energies, yielding a new parameter set called SIRAH-OBAFE (table S2 for parameter values).

The predicted hydration free energies show a much improved agreement with experiment, compared to the previous version of the SIRAH force field, with increased R^2^ values of 0.97 for the new SIRAH-OBAFE parameter set, against values of 0.1 and 0.4 for the SIRAH 1.0 and SIRAH 2.0 sets. Attempts were made to optimise charged side-chains, using free energy gradients, with the necessary correction gradients. While force field parameters able to give improved estimates of the hydration free energies were derived, given the difficulty in this process to avoid an over-fitted model, even with regularization, and the lack of sufficient parameters to improve the hydration free energies in a physically meaningful way, the original charged parameters of the SIRAH force field were retained. The structural stability of proteins has been improved with the use of the new SIRAH-OBAFE force field. RMSD values were reduced by an average of ∼0.25 nm across the protein system tested (Fig. 3), compared to the original SIRAH 1.0 force field, which was used as the starting point in the optimisation procedures.

We believe that the simplification of the physics observed in coarse-grained force fields, such as the SIRAH model, presents a challenge for the reproduction of multiple experimental properties. Limitations in the optimisation methodology are arguably the main cause for this, mainly given by the size of the parameter set that is available to optimise the property of interest; there is insufficient granularity to capture the physics involved in the calculation of hydration free energies for charged and neutral species in a balanced way. The few parameters available in CG models will likely limit the applicability of our proposed optimisation method. To better understand the implications, future studies could be related to the use of a more complex CG protein force field (near atomistic resolution) in the optimisation process, and different scenarios in terms of protein simulations, such as calculating protein potentials of mean force (PMF) for conformational changes and the folding of small peptides. Moreover, a procedure to simultaneously and automatically include PMF data in the ForceBalance parameterisation might bring improvements. Significant and further validation is needed.

The development of new strategies and approaches for force field optimisation is of great interest. In this matter, our new method opens a door for the improvement of contemporary, or new, CG force fields, and it greatly increase the applicability of the CG models in different research areas, such as the study of protein conformational changes, which needs of a correct description of protein-protein, and protein-solute interactions. Furthermore, the parameterisation approach opens a new route to developing CG force fields for other classes of biomolecules such as carbohydrates, nucleic acids, lipids and metabolites, where experimental data may not be as readily available.

## Conflicst of Interest

The authors declare no competing financial interests.

## Acknowledgements

Calculations in this work made use of the Iridis4 supercomputers at the University of Southampton. LPW acknowledges funding support from NIH R01 AI130684-01A1. J.C.D. gratefully acknowledge funding support from CONICYT-BECAS CHILE.

## Supplementary Information

### ForceBalance optimisation procedure

A trust region method approach has been used in the ForceBalance optimisations in this work. In trust region methods, there is a region of search space in which it is assumed the local derivative information is a good approximation of the objective function being minimised. After each optimisation step, the trust radius may be increased or decreased based on the quality *Q* of the steps taken, i.e. the ratio of the objective function change between steps *i* and *i*+1 and the expected change from the local derivative information at step *i*. The following formula is used to adjust the adaptive trust radius after the step is taken:

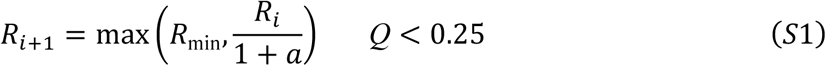

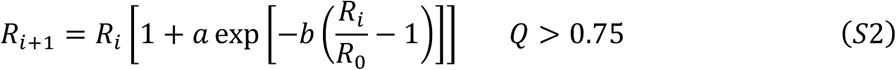

Here *R*_*i*_ is the current trust radius; *R*_*i*+1_ the trust radius at the next iteration; *R*_0_ the default trust radius, was set to 0.1; and *R*_min_ the minimum trust radius, was set to 0.05. The parameter *a*, called “adapt_fac” in ForceBalance, which is related to how much the step size is increased, was set to 1.0; *b*, called “adapt_damp”, that ties down the trust radius, was set to 0.2. The exponential term biases the current trust radius toward the default value, i.e. the trust radius increases by larger factors if the current value is smaller than the default, and vice versa if larger.

### ForceBalance optimisation based on hydration free energy gradients

The optimisation of side-chain analogues and the protein backbone has been made based on atomistic hydration free energies, following 4 stages.

### Stage A: Hydration free energy calculations on AT side-chain analogues, CG side-chains and backbone beads

The interaction energy terms between the solute and solvent are linearly related to the coupling parameter α. With this, the solvation free energies for the side-chain analogues, for the atomistic and coarse-grained systems, were calculated based on a decoupling approximation. That is, interactions between the solute and the solvent were gradually turned off. Our reference state will be our system in solution, and the final state will be the solute in vacuum. The OPLS-AA^1^ and the AMBER-14SB^2^ force fields were used for the atomistic side-chain analogues and the backbone, respectively. In all cases, systems were solvated in a TIP3P^3^ water box. Since we are comparing our calculations with previous studies, especially the ones that give closer results to experiments, we have tried to be consistent with those, hence the choice of different force fields on the optimisation process. As we are taking the best of different worlds to optimise a CG force field, we believe this choice is the most sensible throughout this protocol. The SIRAH protein force field^4^ was used for the CG side-chains and backbone beads, solvated in a WT4^5^ water box. Electrostatic and van der Waals interactions were turned off together. Eleven discrete values of the coupling parameter α were used for the scaling of both CG and AT side-chain analogues potentials (see table 2 and table S1 for details on the analogues used): 0.0, 0.1, 0.2, 0.3, 0.4, 0.5, 0.6, 0.7, 0.8, 0.9 and 1.0, where 0.0 and 1.0 represent the fully on and fully off systems. In the case of N-methylacetamide (NMA), which was used as a representation of the backbone beads, twenty-five values were used: 0.0, 0.5, 0.1, 0.15, 0.2, 0.25, 0.3, 0.35 0.4, 0.45, 0.5, 0.55, 0.6, 0.65, 0.7, 0.75, 0.8, 0.85, 0.87, 0.89, 0.91, 0.93, 0.95, 0.97 and 1.0. The soft-core scaling methods for Lennard-Jones (with α_LJ_ = 0.5) and Coulombic interactions were used to smoothly vary the potentials^6,7^. Simulations were run for 5 ns per window, with a previous equilibration of 1 ns and 5000 iterations of the steepest descent algorithm. All the simulations were run using the NPT ensemble. The Multistate Bennett Acceptance Ratio (MBAR)^8^ was used to compute the free energy difference, which combines data from multiple states. This method is an extension of the well-known Bennett Acceptance Ratio (BAR)^9^, which needs information from two states (in contrast to FEP^10^ or TI^11^, which need information from only one state) in order to compute the free energy difference.

For the AT simulations, a leap-frog stochastic dynamics integrator was used for integration of Newton’s equations of motion with a time-step of 2 fs. Electrostatics interactions were calculated using the PME procedure^12^ with a real-space cut-off of 1.2 nm and a Fourier grid spacing of 0.12 nm. Van der Waals interactions were modelled using the classical Lennard-Jones potential with a cut-off of 1.2 nm. The LINCS algorithm^13^ was applied to constrain all H-bond lengths. AT simulations were run at 1 atm with the Parrinello-Rahman barostat^14^ and at 298.15 K with the Berendsen thermostat^15^.

For the CG simulations, a leap-frog stochastic dynamics integrator was used for integration of Newton’s equations of motion with a time-step of 20 fs. Electrostatic interactions were calculated using the PME procedure with a grid spacing of 0.2 nm. Non-bonded interactions were modelled using the classical Lennard-Jones potential and a Coulombic energy function, with a cut-off of 1.2 nm each. All simulations were run at 1 atm with the Parrinello-Rahman barostat and at 298.15 K with the v-rescale thermostat^16^. All simulations were run with GROMACS v. 2018.2^17^.

### Stage B: Collection of AT <ΔU>α values

<ΔU>_α_ were collected from the AT simulations in stage A, at different α values. For most of the side-chains, α simulations at 0.0 were not used due to the large magnitudes of <ΔU>_α_ values and differences between AA and CG that could not be closely fitted. Val, Cys and Trp were the only exceptions for this case. <ΔU>_α_ values were collected with an in-house Python code created for this purpose, averaging ΔU values for each frame in the trajectories. Table S1 summarise the α values used for each of the simulated side-chains in the ForceBalance optimisation.

### Stage C: Optimisation of SIRAH CG side-chains and backbone

Derivatives of the free energy gradients with respect to the parameters are calculated. These are used to build an objective function, which is a squared sum of the differences between the AA and CG <ΔU>_α_ values. The optimisation was carried out using ForceBalance using the same settings described in the WT4 model development, except for the adapt_fac and adapt_damp parameters, that were set to 0.2 and 0.5 respectively. Only 10 sets of parameters were optimised, 9 of them corresponding to 13 un-charged amino acid side-chains, as some of the side-chains are described by identical parameters, and 1 set corresponding to the backbone beads. In this case, the targets were atomistic free energy gradients at 2 or 3 different α simulation values (table S1). Proline is the only side-chain that has not been optimised given the lack of side-chain analogues, keeping its original parameter values. Only non-bonded parameters were optimised, including van der Waals epsilon (ε) values, and charges, mainly given the parameter sensitivity observed (see below for a discussion on parameter dependence and figure S1). All new optimised parameters are shown in table S2. All the optimisation simulations for the SIRAH beads were run with the optimised WT4-FB model (this work). 100 optimisation cycles were carried out, and the optimal parameters were taken from the lowest value of the objective function. Systems were minimised for 5000 steps using a steepest descent algorithm followed by an NPT equilibration time of 5 ns. Production runs were performed for 10 ns. A leap-frog algorithm was used for integration of Newton’s equations of motion with a time-step of 20 fs. Electrostatic interactions are calculated using the Particle Mesh Ewald method^12^ with a direct cut-off of 1.2 nm and a grid spacing of 0.2 nm. A 1.2 nm cut-off was used for van der Waals interactions. The v-rescale thermostat^16^ and the Parrinello-Rahman barostat^14^ were used to maintain the temperature at 298.15 K and the pressure at 1 atm, respectively. The simulation conditions were consistent with the original SIRAH publication^4^. All simulations were run with GROMACS v. 2018.2^17^. All specific non-bonded pairs, previously set to the original SIRAH force field, between the backbone beads (GC, GN and GO) and water beads (WT) have been removed, and we have set those interactions using Lorentz-Berthelot combing rules.

### Stage D: Re-calculation of CG hydration free energies

The optimised SIRAH-OBAFE force field was used for the re-calculation of the coarse-grained hydration free energies. The same protocol in stage A was used, with some minor differences based on the original publication of the SIRAH protein force field^4^. For all simulation, a leap-frog stochastic dynamics integrator was used for integration of Newton’s equations of motion with a time-step of 20 fs. Electrostatic interactions were calculated using the PME procedure with a grid spacing of 0.2 nm. Non-bonded interactions were modelled using the classical Lennard-Jones potential and a Coulombic energy function, with a cut-off of 1.2 nm each. The LINCS algorithm was applied to constraint all H-bond lengths. All simulations were run at 1 atm with the Parrinello-Rahman barostat^14^ and at 298.15 K with the v-rescale thermostat^16^. All simulations were run with GROMACS v. 2018.2^17^. The new hydration free energies are shown in table 2, and they are compared with hydration free energies calculated from atomistic systems, with the SIRAH 1.0 and the updated SIRAH 2.0 protein force fields.

### Hydration free energies of charged side-chains

Raw hydration free energies (equation 5) have been calculated using a lattice-summation scheme (PME) by decoupling the interactions, electrostatic and van der Waals together, of the ion (side-chain) with the solvent (excluding intramolecular interactions). Eleven lambda values have been used (0.0, 0.1, …, 0.9, 1.0) for all the charged side-chains, using the GROMOS 54A8^18,19^ for atomistic systems, the original SIRAH 1.0, the updated SIRAH 2.0, and SIRAH-OBAFE force fields in GROMACS v.2018.2. The simulation conditions and soft-core potential settings were similar to the ones used in the calculation of hydration free energies for uncharged side-chains (Stage A from the workflow in figure 1). A standard state correction was used with a value equal to 1.9 kcal•mol^-1^ for water, using a density value of 997 kg•m^-3^ (see refs 18, 20, 21). All the reported raw free energies exclude the self-interaction energy.

The main idea behind the necessary correction for the calculation of hydration free energies for charged systems is to approximate the perturbation introduced by these specific errors by the corresponding perturbation within an idealised system (Born model: non-periodic and Coulombic interactions), and see how much the calculated electrostatic potentials deviate from this “real” value. Some of the introduced corrections are summarised (see refs 18, 20, 21 for more details):

A. Approximate representation of the electrostatic interactions (non-Coulombic) which lead to a deviation of the solvent polarization around the ion relative to an idealised Coulombic system, with also incomplete interactions of the ion with the solvent beyond the cut-off. This type A correction is specific for the electrostatic scheme used; it does not apply for lattice-summation schemes (PME), which are Coulombic in the limit of infinite system sizes, but it does apply for cut-off truncation (CT) or reaction field schemes (BM). The type A correction is specific for the electrostatic potential used, and is evaluated using the same potential, but in the idealized context of a macroscopic and non-periodic system. Moreover, it can be sub-divided into corrections A1 and A2 for CT schemes, which apply beyond the cut-off sphere of the ion and within it, respectively.
B. Approximation of the size of the systems (finite), which do not follow a macroscopic regime. This leads to deviations on the solvent polarization, relative to the polarisation of an ideal system (macroscopic). A clear example is the use of a computational box simulated under periodic boundary conditions. This type B correction is applied for the specific electrostatic scheme in the simulation (e.g., LS, CT or BM scheme).
C. Deviation of the solvent generated electric potential at the atomic site of the ion relative to a “correct” electric potential, which is a consequence of the use of an inappropriate summation scheme for the calculation of electrostatic interactions (i.e. P scheme, which stands for summing over individual charges, and a M scheme, which stands for summing over whole solvent molecules), as well as the presence of a constant potential offset. This type C correction is applied for a specific electrostatic scheme and choice of boundary conditions, and can be subdivided in type C_1_ and C_2_ corrections, for improper potential summation and for the potential offset, respectively.
D. Approximate force-field representations, especially related to the wrong dielectric constant for the solvent model used.

Numerical solutions of the Poisson equation are needed to obtain an estimation of the charging free energy in an idealised system that obeys a macroscopic regime (non-periodic with Coulombic electrostatic interactions) and based on the experimental solvent permittivity 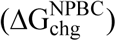. Simulations of a periodic systems with a specific electrostatic scheme and based on the model solvent permittivity are also needed (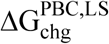 for a periodic boundary condition system using a LS scheme). The sum of corrections A, B and D can be obtained based on these two continuum-electrostatic simulations, as

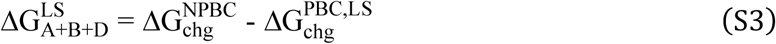

for a LS scheme. The two terms on the right side of equation S1 are charging free energies obtained with the Poisson equation solver from references^22-24^, for non-periodic and periodic systems with Coulombic electrostatic interactions, respectively.

In this work, a relative permittivity of 78.4 for water has been used in the calculation of 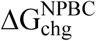 A relative permittivity of 63.84 for the optimised WT4 water model was used, as calculated based in reference 26, in the calculations of 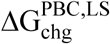. Continuum-electrostatic calculation were done with the GROMOS++ pre-MD and analysis software v.1.4.1^25^ and were based on single structure taken from as the final configuration of the hydration free energy simulations of the charged side-chains. The appropriate boundary conditions and electrostatic scheme were used for each case, with a grid spacing of 0.02 nm and a threshold of 10^−6^ kJ•mol^-1^ for the convergence of the electrostatic free energy.

Type C_1_ correction is required for LS and BM (reaction field) schemes, and corrects the P-summation (atom-based cut-off) implied by these schemes to a proper M-summation (molecule-based cut-off). For a LS scheme, this is given by:

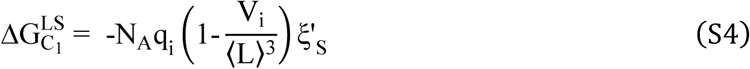

where N_A_ is the Avogadro’s constant, V_i_ is the ionic volume (assumed constant and defined as the change in the volume of the computational box upon insertion of the neutral ion-sized cavity)^18^, L is the length of the edge of the box, q_i_ is the total ionic charge, and ξ’_S_ corresponds to the exclusion potential of the solvent model. For fully rigid models with a single van der Waals interaction site, this last term has been usually calculated based on the quadrupole moment trace of the solvent model. For more complex solvent models, different methods have been derived for the calculations of their exclusion potentials^26^. In this work, we have employed method IV from reference 26, which relies on the comparison of the raw potentials within a cavity using two different electrostatic schemes, assuming that the corrected potentials are equal. For this, we have used a cut-off truncation (CM) and reaction field schemes (BM). The difference in the raw potentials are related to ξ’_S_ as:

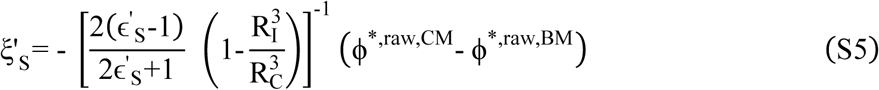

where R_I_ is the effective ionic radius, R_C_ is the cut-off, ϕ^*,raw,CM^ and ϕ^*,raw,BM^ are the raw electrostatic potentials within an uncharged cavity of the size of a CG sodium ion, and ϵ’_S_ corresponds to the dielectric permittivity of the solvent model, which has been calculated based on the methodology used in reference 26. Simulations of an un-charged sodium ion solvated in the optimised WT4-FB model were run for 1 ns using a BM scheme, with a reaction field permittivity ϵ_RF_ equal to 80. Electrostatic potentials at the cavity were obtained for both CM and BM schemes based on the electrostatic interaction of the cavity with the solvent within the cut-off R_C_, using an *in-house* Python script created for this purpose. Simulation settings were similar to the previous one used in this work. The dielectric permittivity calculated here differs with the value previously reported in table 1, but is within the error. Moreover, it has been reported that dielectric permittivities calculated using a reaction-field scheme are more sensitive to the choice of simulation parameters such as the non-bonded cut-off. Given this, the lack of agreement is not unexpected, but as a matter of consistency with previous studies^26^, we decided to use the dielectric permittivity calculated in this section for the evaluation of the exclusion potential. Moreover, the dielectric permittivity for the WT4 water model calculated in this section is similar to the one calculated by Reif et.al^26^ with a reported value of 66.7 using the SPC model.

Type C_2_ corrections correct for the presence of an interfacial potential at the ion surface. This term is proportional to the ratio of the ionic volume to the box volume. With this, its magnitude is very small for the systems used in this work, and has been neglected in the calculation of the corrected hydration free energies.

### Optimisation of charged side-chains

All the optimisation simulations for the SIRAH beads were run with the optimised WT4-FB model (this work). 100 optimisation cycles were run. Systems were minimised for 5000 steps using a steepest descent algorithm followed by an NPT equilibration time of 5 ns. Production runs were performed for 10 ns. A leap-frog algorithm was used for integration of Newton’s equations of motion with a time-step of 20 fs. Electrostatic interactions are calculated using the Particle Mesh Ewald method^12^ with a direct cut-off of 1.2 nm and a grid spacing of 0.2 nm. A 1.2 nm cutoff was used for van der Waals interactions. The v-rescale thermostat^16^ and the Parrinello-Rahman barostat^14^ were used to maintain the temperature at 298.15 K and the pressure at 1 atm, respectively. The simulation conditions were consistent with the original SIRAH publication^4^. All simulations were run with GROMACS v. 2018.2^17^.

For the PMF calculations, the distance between the centre of mass of the BCG bead of Glu, and the BCE bead of Lys and BCZ bead of Arg, were used as collective variables. A total of 78 windows have been used for umbrella sampling for both cases with distances spanning from 0.38 nm to 1.8 nm, using a spring constant of 5000 kJ·mol^-1^. Simulations settings were similar to the previous one used: a leap-frog stochastic dynamics integrator was used for integration of Newton’s equations of motion with a time-step of 20 fs. Electrostatic interactions were calculated using the PME procedure with a grid spacing of 0.2 nm. Non-bonded interactions were modelled using the classical Lennard-Jones potential and a Coulombic energy function, with a cut-off of 1.2 nm each. The LINCS algorithm was applied to constraint all H-bond lengths. All simulations were run with GROMACS v. 2018.2 at 1 atm with the Parrinello-Rahman barostat and at 298.15 K with the v-rescale thermostat, and were preceded by the corresponding minimisation and NPT equilibration.

**Table S1.** α simulation values used for the collection of <ΔU>_α_ values, that correspond to the targets in the optimisation of the CG beads in ForceBalance. Atomistic analogues used are shown in parenthesis.

| Side-chain | $\alpha$ values |
| --- | --- |
| Asn (acetamide) | 0.1, 0.2, 0.5 |
| Cys (methanethiol) | 0.0, 0.2, 0.4 |
| His (methylimidazole) | 0.1, 0.2, 0.4 |
| Met (methyl-ethylsulfide) | 0.1, 0.2, 0.4 |
| Phe (toluene) | 0.5, 0.6 |
| Ser (methanol) | 0.1, 0.2, 0.5 |
| Trp (methyldole) | 0.0, 0.4, 0.5 |
| Tyr (*p*-cresol) | 0.1, 0.4, 0.5 |
| Val (propane) | 0.0, 0.5 |
| Backbone (N-methylacetamide) | 0.1, 0.3 |

### Parameter dependence

Initially, a screening test was performed to evaluate the parameter dependence of <ΔU>_α_ with respect to the force field parameters, i.e. to evaluate the changes in <ΔU>_α_ based on changes in the force field parameters. For some of the cases (Ser, Asn and Val), both van der Waals sigma (σ) and epsilon (ε) values were optimised in a first attempt. Based on the parameter dependence observed in figure S2 for the case of Val, the <ΔU>_α_ values do not significantly change within a sensible range of van der Waals σ values. On the contrary, an important parameter dependence is shown with respect to the van der Waals ε values (i.e. big changes in <ΔU>_α_ are observed when changes in the parameters are performed). In figure S2, the van der Waals parameters are plotted in the form of internal optimisation variables in ForceBalance (“mathematical parameters”), which are related to the physical parameters (i.e. the parameters that are actually printed in the force field file) as a shifted displacement form the original value:

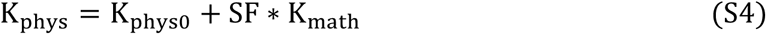

where K_phys_ corresponds to the parameter that is used in the simulation after the optimisation process, K_phys0_ is the initial parameter before the optimisation, SF is the scaling factor and K_math_ the mathematical value used in the optimisation process. Finally, only van der Waals ε values and charges were optimised given the parameter sensitivity that exists (see table S2 and figure S2).

**Figure S1.**
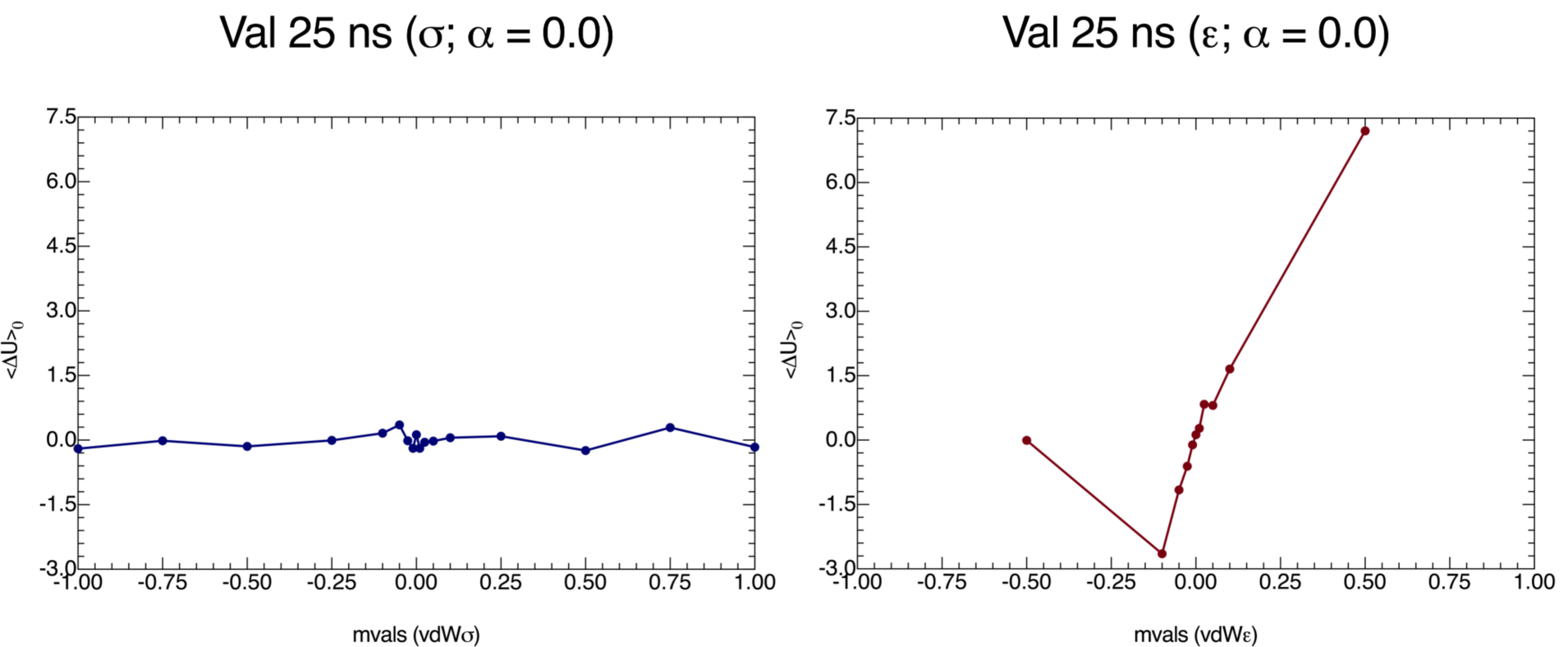
Parameter dependence for Val. Changes of <ΔU>_0_ (in units of kcal·mol^-1^) with respect to the van der Waals (vdW) σ and ε values are shown (left and right panel, respectively). Simulations were run at α = 0.0 (fully on solute) for 25 ns. The simulation conditions were the same as the ones used for the side-chain optimisations (see stage A). Van der Waals values are plotted as mathematical values (mvals).

### Atomistic gradient choice

It is important to note that for most cases, AT gradients at α=0 were too high to be fitted by ForceBalance due to the large magnitude of the gradient and the large difference between AT and CG. Inclusion of the α=0.0 point would have introduced a very large contribution to the objective function and worsened the quality of fit of all the other α points. We are assuming that the gradients should behave in a similar manner between the all-atom and coarse-grained systems, but this might not be the case. Using the free energy gradients as a proxy for the free energies, instead of the free energy itself, relies on the assumption that 1) if one of the free energy gradients is correct, we expect a better performance across the whole range of α values, and 2) coarse-grained and atomistic systems should have similar free energy gradients. Neither of these is necessarily true.

### The methionine case

The optimisation of methionine is an example where our method has worked (Fig. S2), finding a minimum, i.e. the optimal set of parameters to minimise the objective function. A manual search of 441 parameter combinations shown in figure S2 led us to similar results to those obtained for the full optimisation of methionine, with values of vdWσ = 0.49 nm, vdWε = 4.56 kJ·mol^-1^, and vdWσ = 0.48 nm and vdWε = 4.22 kJ·mol^-1^, respectively. Figure S3 shows the free energy gradients for the atomistic and coarse-grained methionine side-chain. The overall shape of the profile is maintained, but differences exists in the magnitude of the gradients. This may account for the differences observed for the calculated HFEs. Fortuitously, the optimised parameters led to better agreement with experimental hydration free energies.

In the case of phenylalanine, the optimised SIRAF-OBAFE parameters performed worse compared to the original SIRAH force fields, with values of 0.50 ± 0.05 kcal·mol^-1^ for the SIRAH 1.0 force field, 0.57 ± 0.05 kcal·mol^-1^ for the SIRAH 2.0 force field, vs. 1.12 ± 0.06 kcal·mol^-1^ for the optimised SIRAH-OBAFE. We believe this is mainly due the complexity on the free energy gradient profile for this residue. Moreover, later optimisation runs of side-chains that share the A2C bead-types with phenylalanine (such as His, Tyr, and Trp) were performed using this parameter fixed to its original value.

**Figure S2.**
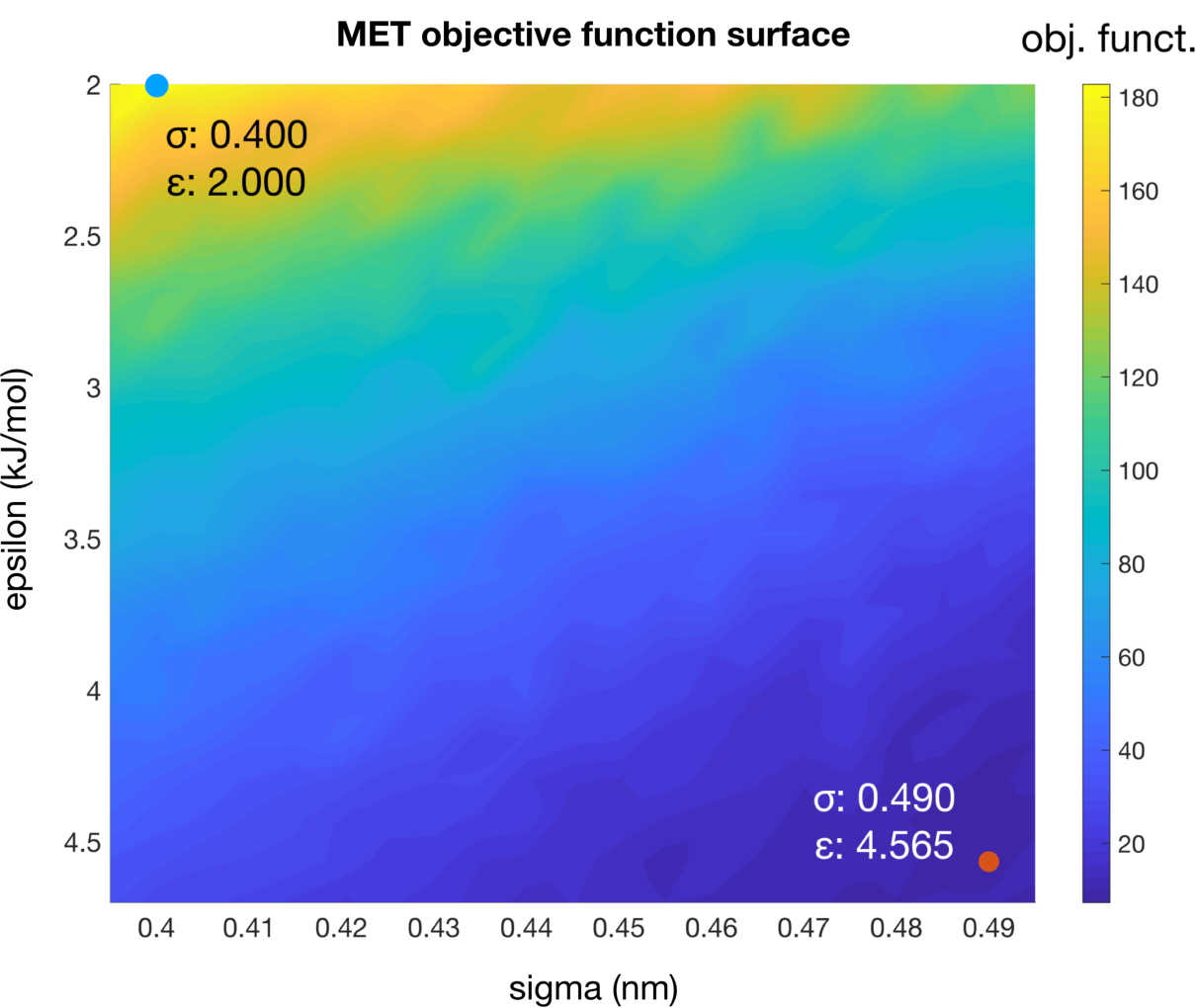
Methionine objective function surface. 441 combinations (21 x 21) of vdWσ and vdWε simulations were performed, and single calculations of the objective function were extracted and plotted. The maximum and minimum values for the objective function are shown as blue and red dots, respectively, and for each of these the parameter combination is shown.

**Figure S3.**
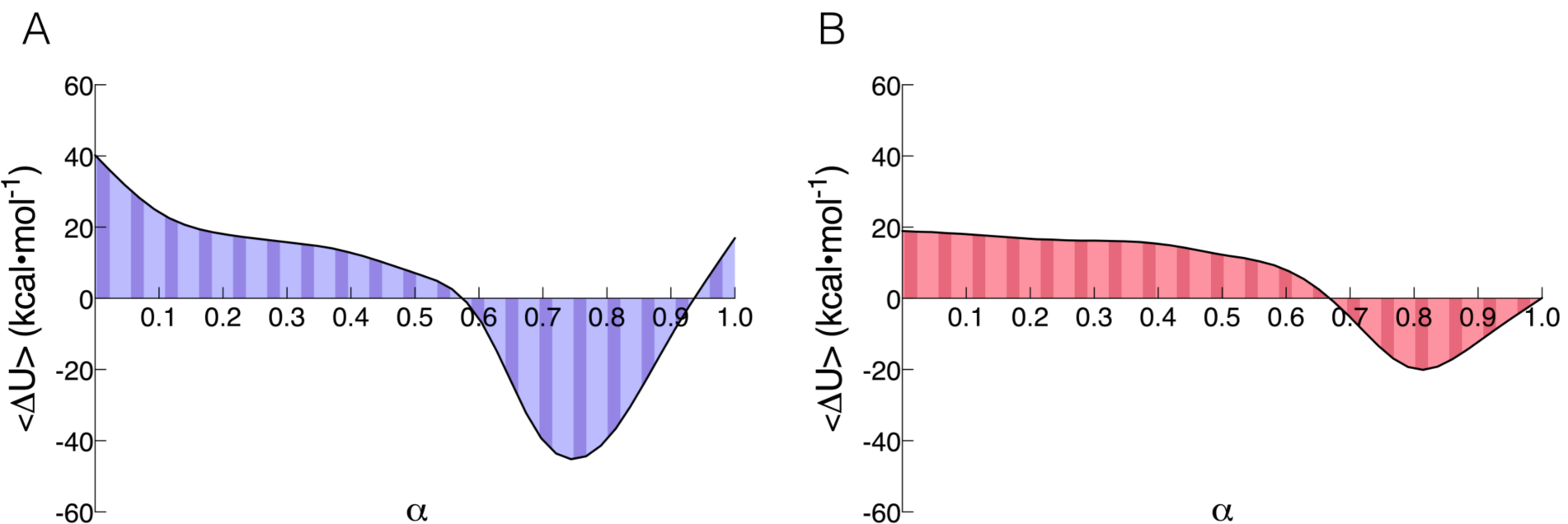
Free energy gradients for methionine. (A) atomistic free energy gradients and (B) coarse-grained free energy gradients. The results represent 11 α simulations with average <ΔU>_α_ values for each of those simulations shown.

**Figure S4.**
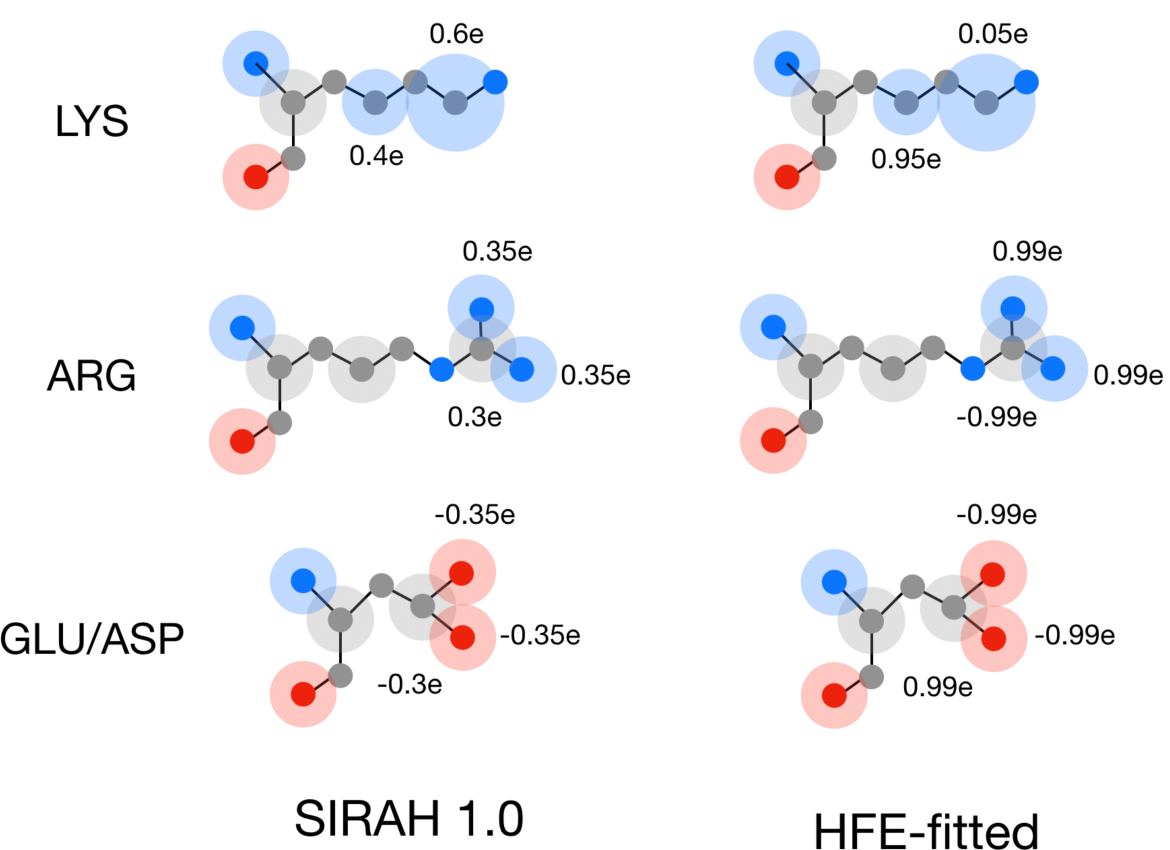
HFE-fitted charge values for the optimised charged side-chain. Schematic representation of the three optimised charged side-chains (Lys, Arg and Glu/Asp), for the original SIRAH 1.0 and the HFE-fitted parameter set (after the initial optimisation).

**Figure S5.**
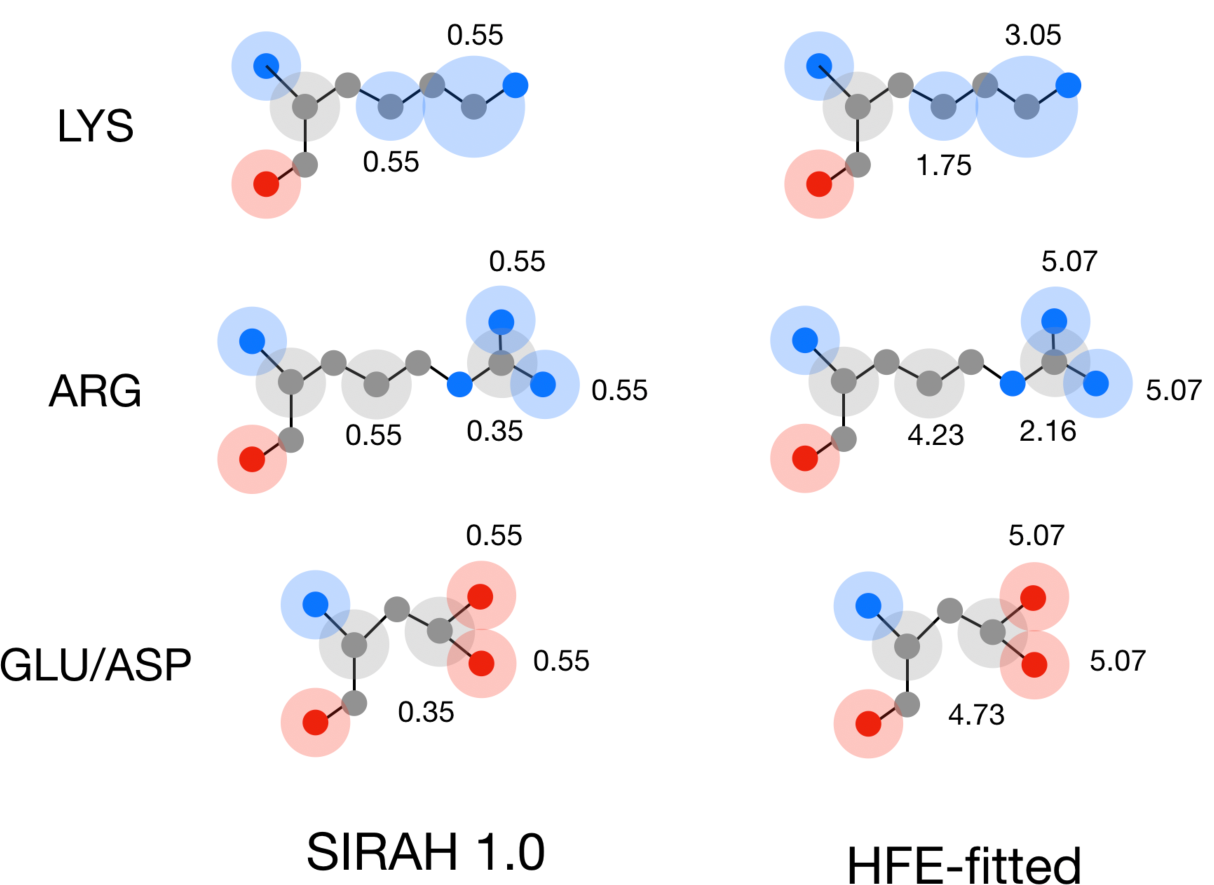
HFE-fitted VDWε values for the optimised charged side-chain. Schematic representation of the three optimised charged side-chains (Lys, Arg and Glu/Asp), for the original SIRAH 1.0 and the HFE-fitted parameter set (after the initial optimisation). All values are in units of kJ·mol^-1^.

**Figure S6.**
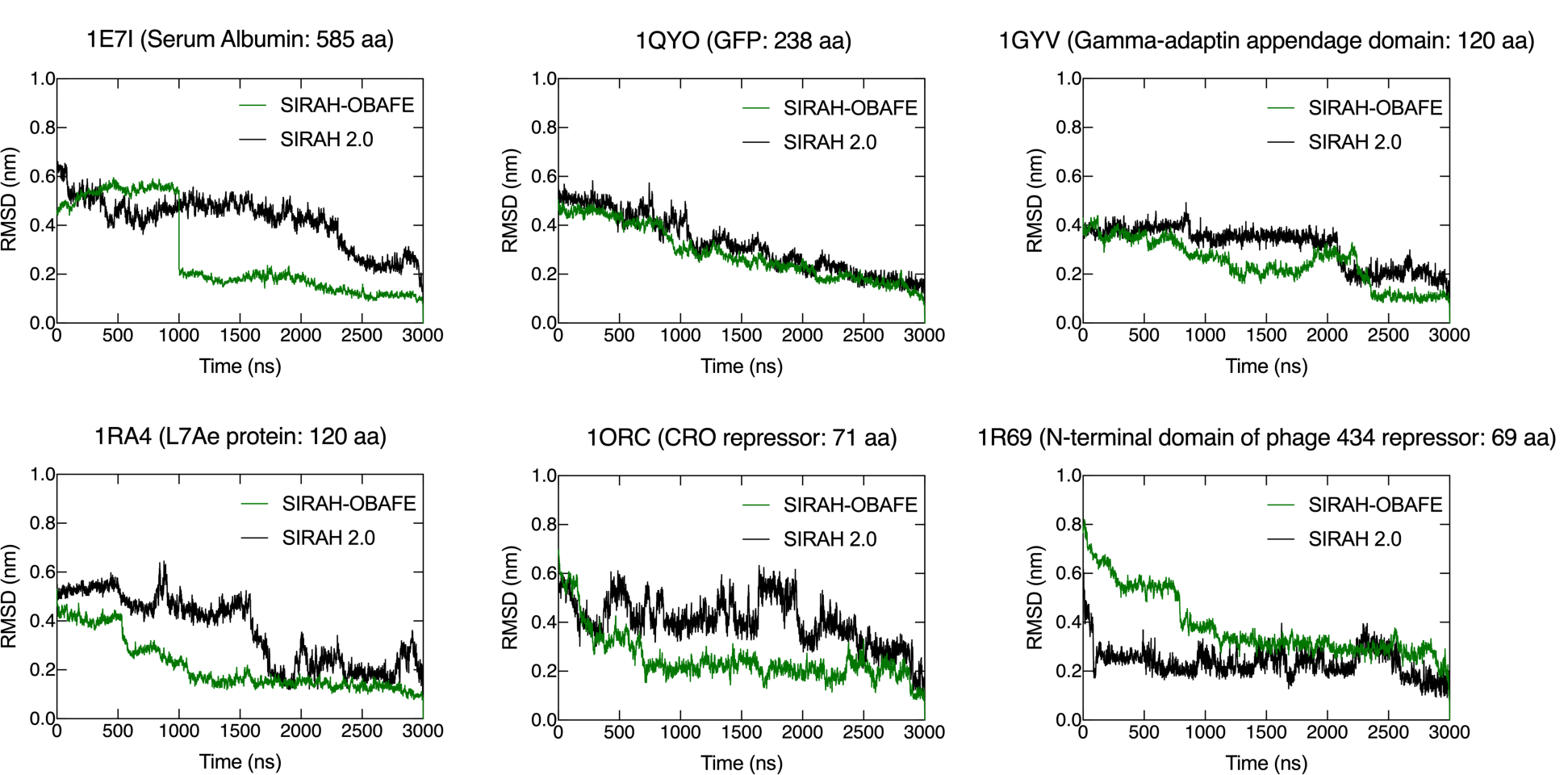
RMSD times series against the last frame. RMSD trajectory analysis is shown as a time series comparison with respect to the Cα carbons of the CG representation to the last frame of the trajectory for (A) Serum albumin, (B) GFP protein, (C) Gamma-adaptin domain, (D) L7Ae Archeal ribosomal protein, (E) CRO repressor and (F) the N-terminal domain of phage 434 repressor. PDB codes are shown in the figure titles. Simulations were run using the SIRAH 2.0 (black) and SIRAH-OBAFE (green) force fields.

**Table S2.** Optimised parameters for the SIRAH-OBAFE and WT4-FB force fields

| | Bead type<sup>a</sup> | VDWσ (nm) | VDWε (kJ·mol<sup>-1</sup>) | Charge (*e*) |
| --- | --- | --- | --- | --- |
| Side-chain beads (SIRAH-OBAFE) |  |  |  |  |
| Asn/Gln | P3Cn/q | SaO<sup>b</sup> | 3.5217E-01 | 0.00 |
|  | P5N | SaO | 5.5453E-01 | 5.9527E-01 |
|  | P4O | SaO | 5.547E-01 | -5.9527E-01 |
| Cys | P1S | SaO | 1.0547E+00 | -6.0817E-01 |
|  | P2P | SaO | 2.2622E-01 | 6.0817E-01 |
| His (epsilon protonated) | A2C | SaO | SaO | 0.00 |
|  | A5E | SaO | 1.7084E+00 | 5.0449E-01 |
|  | A5D | SaO | 1.7023E+00 | -5.0449E-01 |
| Met | Y3Sm | SaO | 4.7181E+00 | 0.00 |
| Phe | A2C | SaO | SaO | 0.00 |
|  | A1C | SaO | SaO | 0.00 |
| Ser/Thr | P1O | SaO | 4.4658E-01 | -9.1874E-01 |
|  | P2P | SaO | 2.2622E-01 | 9.1874E-01 |
| Trp | A2C | SaO | SaO | 0.00 |
|  | A7N | SaO | 6.9916E-01 | -3.5323E-01 |
|  | A8P | SaO | 1.5469E+00 | 3.5323E-01 |
|  | A1Cw | SaO | 3.1449E+00 | 0.00 |
| Tyr | A2C | SaO | SaO | 0.00 |
|  | A4O | SaO | 2.0418E+00 | -3.5107E-01 |
|  | A3P | SaO | 2.0491E+00 | 3.5107E-01 |
| Val/Leu/Ile | Y4Cv/Y1C | SaO | 5.0887E-01 | 0.00 |
| Backbone | GC | SaO | 5.5058E-01 | 4.2176E-01 |
|  | GO | SaO | 5.2511E-01 | -6.7336E-01 |
|  | GN | SaO | 5.5058E-01 | 2.5161E-01 |
| Arg | C2Cr | SaO | SaO | SaO |
|  | C3Cr | SaO | SaO | SaO |
|  | C5N | SaO | SaO | SaO |
| Lys | C1Ck | SaO | SaO | SaO |
|  | C7Nk | SaO | SaO | SaO |
| Asp/Glu | C4Ce/d | SaO | SaO | SaO |
|  | C6O | SaO | SaO | SaO |
| **WT4-FB** |  |  |  |  |
|  | WN1 | 4.2474E-01 | 7.6717E-01 | -2.6730E-01 |
|  | WN2 | 4.2474E-01 | 7.6717E-01 | -5.6223E-01 |
|  | WP1 | 4.2474E-01 | 7.6717E-01 | 2.6730E-01 |
|  | WP2 | 4.2474E-01 | 7.6717E-01 | 5.6223E-01 |

<sup>a</sup> Bead types taken from the original SIRAH publication<sup>4</sup>.
<sup>b</sup> SaO, same as original, taken from the SIRAH protein force field publication<sup>4</sup>.

**Table S3.** Hydration free energies of neutral side-chains and backbone using the OPLS-AA, AMBER-14SB, SIRAH 1.0, SIRAH 2.0 and SIRAH-OBAFE force fields^a,b^

|  | **Expt.** | **OPLS-AA<sup>c</sup>** | **SIRAH 1.0<sup>c</sup>** | **SIRAH 2.0<sup>c</sup>** | **SIRAH-OBFAFE<sup>c</sup>** |
| --- | --- | --- | --- | --- | --- |
| Backbone (NMA) | 10.1 | 7.40 ± 0.04<br>(AMBER-14SB) | -1.73 ± 0.07 | -0.16 ± 0.05 | 10.91 ± 0.05 |
| Val (propane) | -1.99 | -2.45 ± 0.06 | -0.02 ± 0.01 | -0.18 ± 0.01 | -2.26 ± 0.03 |
| Leu (isobutane) | -2.28 | -2.69 ± 0.10 | -0.02 ± 0.01 | -0.18 ± 0.01 | -2.26 ± 0.03 |
| Ile (butane) | -2.15 | -2.59 ± 0.08 | -0.02 ± 0.01 | -0.18 ± 0.01 | -2.26 ± 0.03 |
| Ser (methanol) | 5.06 | 4.44 ± 0.01 | -1.87 ± 0.04 | 0.10 ± 0.09 | 5.26 ± 0.10 |
| Thr (ethanol) | 4.88 | 4.12 ± 0.11 | -1.87 ± 0.04 | -0.40 ± 0.01 | 5.26 ± 0.10 |
| Cys (methanethiol) | 1.24 | 0.39 ± 0.02 | -1.78 ± 0.03 | -0.71 ± 0.01 | 0.92 ± 0.07 |
| Met (methyl-ethylsulfide) | 1.48 | 0.06 ± 0.01 | -0.03 ± 0.02 | -0.01 ± 0.03 | 1.36 ± 0.02 |
| Asn (acetamide) | 9.68 | 8.46 ± 0.02 | 2.87 ± 0.07 | 2.85 ± 0.04 | 8.12 ± 0.05 |
| Gln (propionamide) | 9.38 | 8.36 ± 0.04 | 2.87 ± 0.07 | 2.86 ± 0.04 | 8.12 ± 0.05 |
| Phe (toluene) | 0.76 | 0.40 ± 0.04 | 0.50 ± 0.05 | 0.57 ± 0.05 | 1.12 ± 0.06 |
| Tyr (*p*-cresol) | 6.11 | 4.61 ± 0.13 | 0.70 ± 0.06 | 0.67 ± 0.02 | 5.10 ± 0.04 |
| His (methylimidazole) | 10.27 | 7.70 ± 0.06 | 1.47 ± 0.04 | 1.24 ± 0.02 | 8.46 ± 0.08 |
| Trp (methyldole) | 5.88 | 5.55 ± 0.22 | -0.47 ± 0.09 | 1.52 ± 0.02 | 4.51 ± 0.04 |
| **MUE<sup>b</sup>** |  | **1.04** | **5.03** | **4.45** | **0.68** |
| **MSE<sup>b</sup>** |  | **-1.04** | **-4.13** | **-3.61** | **-0.43** |
| **R<sup>2</sup>** |  | **0.98** | **0.10** | **0.40** | **0.97** |

<sup>a</sup> Values are in the units of kcal·mol<sup>-1</sup>. Experimental values were obtained from reference 27. OPLS-AA and AMBER-14SB values were re-calculated using the corresponding side-chain analogues listed in parenthesis, based on reference <sup>26</sup>.
<sup>b</sup> Mean signed error (MSE), mean unsigned error (MUE) and determination coefficient (R<sup>2</sup>).
<sup>c</sup> Error bars modelled as standard errors across three repeat simulations.

## Notes

### Competing Interest Statement

The authors have declared no competing interest.

